# Decentralized social media for environmental research

**DOI:** 10.64898/2026.11.06.764567

**Authors:** Abdesslam Chai-allah, Andrea Ghermandi, Nathan Fox, Derek Van Berkel

## Abstract

Social media data are reshaping environmental research. However, recent ownership and policy changes on centralized platforms like Twitter/X have motivated many users to seek new alternatives while simultaneously accelerating data restrictions for scientists. Decentralized social media, particularly Bluesky and Mastodon, are emerging as alternatives for users seeking greater control over their data and as a more stable and sustainable source of data for researchers. This rise may be instrumental in revitalizing the role of social media in environmental research. This study aims to provide a first-of-its-kind analysis of the potential of these new sources for environmental research, demonstrating their potential usefulness through the examples of climate change discourses and people’s appreciation of wildlife. We find an increased adoption of Mastodon and Bluesky, with high linguistic diversity, even if dominated by English. Wildlife image content reveals broad taxonomic diversity, with plants, birds, and insects dominating the records. We find this pattern broadly consistent with that observed on the popular citizen science platform iNaturalist. Climate change discourse is found to broadly reflect the scientific consensus on both platforms, with contrarian claims remaining uncommon. This suggests that researchers may benefit from less polarized opinions and misinformation, arising from interactions that are less driven by engagement maximizing algorithms implemented in other platforms. Our study illustrates the promise of decentralized social media as a new data source for environmental scientists and paves the way for their broader use in environmental research.

## 1. Introduction

Researchers in environmental sciences are increasingly utilizing social media (SM) as a rich data source to analyze public attitudes, behaviors, and perspectives. For instance, SM data have enhanced the monitoring of visitation to protected areas, the valuation of ecosystem services and scenic landscapes, the analysis of climate change communication, natural disaster monitoring, public opinion on renewable energy, and more (Williams et al. 2015; Van Zanten et al. 2016; Moore and Obradovich 2020; Falkenberg et al. 2022; Jeong et al. 2023). However, recent years have seen growing restrictions on data access to some of the largest and globally important platforms, including Twitter/X, following Elon Musk’s takeover of the platform, Meta platforms (Facebook, Instagram), and TikTok. Such developments sharply highlighted the precarious nature of these data sources and the risks involved in reliance on them for long-term monitoring and decision-making. While many researchers have recognized the importance of fostering strong academia-industry collaborations with SM platforms to maintain access to these data, there have also been calls to broaden approaches and explore alternative platforms to make data access more diversified and resilient to changes in individual platforms (Ghermandi et al. 2023). The engagement of leading SM platforms with data sharing for academic research remains limited (e.g., Meta Content Library access through university-led platforms such as SOMAR; https://www.icpsr.umich.edu/sites/somar/meta-content-library), it often requires a complex and lengthy approval procedure that limits their usefulness, and remains open to the risk of being abruptly discontinued from the top of their corporate structure (e.g., Twitter academic research product track).

In this context, emerging SM platforms that rely on federated and decentralized networks present opportunities to fill such gaps while being less exposed to the risks of traditional centralized platforms. This new ecosystem includes multiple SM platforms with similar or analogous affordances to popular corporate SM and with growing, albeit still relatively small, user bases (Fig 1). To date, however, little research has explored these emerging SM platforms in the context of environmental and sustainability science (Schiffman 2025; Zheng et al. 2025; Hanny et al. 2026).

**Figure 1.**
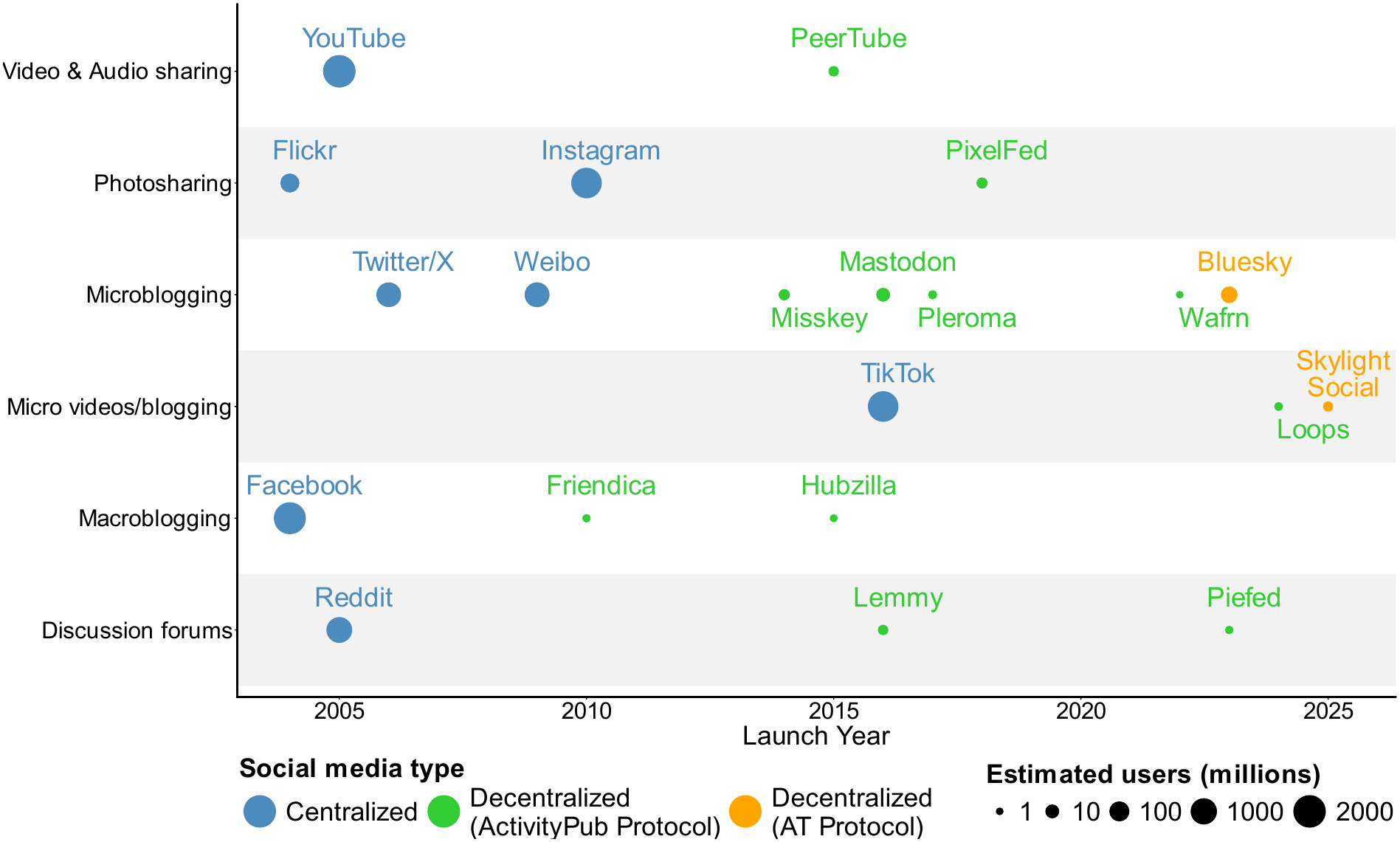
Social Media ecosystem: type, purpose, launch date and total users. For centralized platforms, user counts are from April 2025 and were obtained from DataReportal (https://datareportal.com/social-media-users). User data for Weibo are from the end of 2022 (Ma & Zheng 2025). For decentralized ActivityPub platforms, user counts represent totals as of June 2026 and were obtained from FediDB (https://fedidb.com/). User data for Bluesky and Mastodon were collected directly through their respective APIs.

Decentralized platforms are distinctive from traditional SM platforms, using decentralized or federated models for hosting content, some of which are collectively known under the umbrella of the “fediverse”. “Federated” and “decentralized” are often used interchangeably, but these terms describe different aspects of network architecture (Roscam Abbing et al. 2023). Both concepts imply a departure from centralized control and may enable greater user stewardship and participatory governance, yet the boundary between them is often unclear and inconsistently defined in the literature. Two decentralized SM platforms that have seen rapid growth in interest and user numbers in recent years are Mastodon and Bluesky (Zulli et al. 2020; Failla and Rossetti 2024). Mastodon is both federated and decentralized: it operates through a large number of independently managed servers that distribute hosting and management of user data, communities, and moderation (decentralization), while communicating with one another via interoperable protocols, primarily ActivityPub (federation). In contrast, while still decentralizing user data and identity, Bluesky’s architectural model still allows a degree of centralized influence through application-level services, including content moderation and curation, and the user experience is less dependent on the specific server to which the user belongs. While federated networks are always decentralized, not all decentralized networks are federated (Roscam Abbing et al. 2023). Decentralized platforms span thus a spectrum of different models, rather than representing a simple binary shift from centralized alternatives (Ciriello et al. 2026).

Decentralized networks contrast with traditional, commercially owned centralized SM platforms (e.g., Instagram and Facebook) by shifting control from a single governing organization to a distributed network of users and communities (La Cava et al. 2022). The subset of networks using a federated model (e.g., ActivityPub) creates a diversity of community ownership as each server is managed independently, allowing users to join the server(s) that are best aligned with their preferences or even setting up their own servers while still interacting with users across different servers, both within and beyond their chosen platform, as long as the servers share the same underlying protocol (Zignani et al. 2018; La Cava et al. 2022; Wahner et al. 2024). For example, Mastodon (a microblogging platform), PixelFed (a photo-sharing platform), and PeerTube (a video-sharing platform) rely on the ActivityPub protocol, enabling interoperability and cross-platform interaction for their users.

While the decentralized approach of “fediverse” platforms is built on a system where independent servers store and manage the data of their users, the data remains owned by the server hosting it and is ultimately controlled by the server admins, rather than by the users themselves. Bluesky, through the AT Protocol, is based on a new approach for decentralizing SM. The key innovation of the AT protocol is the introduction of 1) a Personal Data Server for each user, allowing them to store their data independently, and 2) a Decentralized User Identifier, allowing users to create their own cross-platform identifier. The platform built atop the AT protocol (e.g., Bluesky) plays the role of an aggregator that collects data from all the personal data servers in the network, processes it, and displays it to the user via the app (Failla and Rossetti 2024; Kleppmann et al. 2024).

This diversity, even across countries, and community-driven control within decentralized and federated SM networks, presents intriguing possibilities for researchers aiming to surface information less affected by the algorithmic biases and monetization pressures that drive engagement in centralized platforms through polarizing or extreme content (Kim and Dennis 2019; Kitchens et al. 2020). Mainstream platforms have designed algorithms to maximize user engagement by promoting polarizing or extreme content that often amplifies misinformation and divisive rhetoric (Johnson et al. 2019; Avalle et al. 2024). These algorithms are not only shaped by “monetization pressures” but may also be influenced to curry political favor, subtly steering public opinion, or favoring certain narratives (Guess et al. 2023; Avalle et al. 2024). In contrast, decentralized platforms operate without a central authority, allowing communities a much greater control on moderation rules (La Cava et al. 2022; Wahner et al. 2024). This approach may allow for more diverse and inclusive content environments, offering a potential counterbalance to the potential bias in mainstream systems. Moreover, community-driven moderation is likely to align more closely with real-world interactions, where social norms regulate behavior, potentially creating a more authentic digital ecosystem in which content reflects real-world attitudes and opinions. Even in a role of complementary, rather than alternative, sources of data to mainstream platforms, integrating data from such novel platforms into scientific analyses may play an important role in mitigating the measurement and representation biases of studies relying on data from one single SM platform, which remain the vast majority (Ghermandi et al. 2026a). This potential role is further supported by the influence of platform affordances and social norms on who shares content and what they share, with the implication that a larger user base does not necessarily ensure a more accurate representation of user preferences regarding human–nature interactions (Ghermandi et al. 2026b).

The emergence of decentralized SM may thus present new opportunities to collect diverse societal perspectives and large, geographically distributed datasets, and sustainable data access without relying on the governance and algorithms of commercially owned platforms. At the same time, important questions remain regarding their utility, representativeness, and potential biases. What are the growth patterns of their user bases and are they sufficiently sustained over time, or will they remain niche platforms that provide little information that may be of use to environmental researchers? How representative are data derived from them, considering the concern that users may be disproportionately drawn from technologically engaged communities, limiting demographic diversity and fostering ideologically homogeneous communities?

In this study, we provide a first exploratory, large-scale assessment of the potential value of decentralized SM platforms as data sources for environmental research. We focus on two leading decentralized platforms—Mastodon and Bluesky—to characterize their user growth patterns and linguistic diversity, and to provide a comparative quantitative and qualitative assessment of the content available on both platforms pertaining to two important environmental research domains, public discourse on climate change and wildlife photography. More specifically, we assess the prevalence of climate change denial and the relative distribution of different taxonomic groups across the two platforms and compare them to those of other leading platforms. We build on the results to discuss the potential of decentralized SM as complementary and more sustainable sources of data for environmental researchers.

## 2. Materials and Methods

### 2.1. Mapping decentralized networks within the social media ecosystem

To better understand the emergence of decentralized social media, we conducted an inventory of platforms built on the two decentralized protocols ActivityPub and AT Protocol. For each platform, we collected information on year of launch, user base, and platform type (e.g., microblogging, photo-sharing, or video-sharing). We limited our analysis to platforms with at least 5,000 users as of June 2026 in order to focus on platforms that have achieved a sufficient level of adoption and operational sustainability. We then compared these platforms with the corresponding platform types in the centralized social media ecosystem.

For ActivityPub-based platforms, we relied on FediDB (https://fedidb.com), which provides data and analytics on the growing ActivityPub ecosystem. For AT Protocol-based platforms, we used ATProto Alternatives (https://atprotoalternatives.com), which provides descriptions of the decentralized alternatives to centralized social media platforms built on the AT Protocol. Finally, we consulted the official website of each platform to verify and validate the information obtained from these sources.

### 2.2. User growth and language diversity in the leading decentralized networks

To evaluate user growth and activity on Mastodon and Bluesky, the two leading platforms within the decentralized social media ecosystem, the total number of accounts and weekly active users were used as indicators of the platforms’ adoption and sustained use. Monthly user growth (total users) and weekly active users were analyzed from July 2023 to June 2026. For Mastodon, we used the statistics API that provides access to the daily evolution of the platform, including the total number of accounts, active users, and server count (https://api.joinmastodon.org/statistics); weekly user activity was estimated by averaging Mastodon’s daily active users over each week. For Bluesky, we used the *com.atproto.sync.subscribeRepos* firehose of the Bluesky API to track account-creation events and derive the total number of registered accounts (https://atproto.com/specs/sync), while weekly active users were calculated as the number of unique users who generated at least one post that week, extracted from Bluesky activity statistics (https://blue.mackuba.eu/stats/).

To evaluate the linguistic diversity on both platforms, we randomly sampled 1 million posts from each platform in September 2024, where the language was specified. On both platforms, users specify their preferred posting language, and the sampled posts are automatically tagged with corresponding language metadata.

### 2.3. Climate change and wildlife posts

To evaluate the potential of data from Mastodon and Bluesky in advancing environmental research, we conducted extensive data collection on both platforms, focusing on two key topics: climate change and wildlife observation. Mastodon and Bluesky enable users to share various types of media, including images and text posts. These posts, along with their associated metadata, are comparable to those from other microblogging social media platforms currently used in environmental research such as Twitter/X. The data was retrieved in September 2024 using the Mastodon and Bluesky APIs. For Mastodon, we used the *v2/search* API endpoint, which allows for textual queries through all public Mastodon instances (https://docs.joinmastodon.org/client/intro/). For Bluesky, we collected the data using the *app.bsky.feed.searchPosts* API endpoint. The historical search timeframe was set from January 1, 2023, to August 31, 2024.

To identify relevant posts for each topic, we performed a keyword search using a list of topic-specific keywords in five languages: English, French, German, Spanish, and Portuguese (the complete list can be found in Table S1 in supplementary material). As a result, we identified 35,545 posts related to climate change that were shared by 12,374 users on Bluesky, and 40,846 posts shared by 5761 users on Mastodon. Regarding wildlife, 169,943 posts were shared by 57,648 users on Bluesky, compared to 131,230 posts shared by 21,312 users on Mastodon. The collected data fields include the original text content of Bluesky and Mastodon posts, URLs for any attached media, publishing time, posting languages, and user ID.

We filtered the collected datasets using two complementary procedures. The first consisted of filtering steps applied uniformly to both climate change and wildlife observation datasets, while the second comprised criteria adapted to the target elements of each dataset (text posts for climate change and images for wildlife observation). The shared filtering procedure first involved detecting and removing potential bots, defined as accounts that posted more than 50 times per day. In Mastodon, we additionally excluded content from two servers (@botsin.space and @mastodon.bot), which are primarily used by users to run their own bots on the platform. For the wildlife dataset, we filtered the remaining posts to include only those with attached media (images, audio, or video), retaining all image media when included. Since individual posts could contain multiple attachments, each media file was collected. We then retained only images, removed all audio and video files, and eliminated duplicates based on identical image URLs. After filtering, our image dataset consisted of 82,662 images from Bluesky and 65,623 from Mastodon. For the climate change dataset, we removed duplicate posts based on the original text from the same user on the same day, keeping only one post. We then pre-processed the original text for further textual analysis by removing all URLs, @mentions, emojis, and hashtags, punctuation marks, and converting all characters to lowercase. After filtering and preprocessing, our final text dataset consisted of 32,142 posts from Bluesky and 34,940 from Mastodon. Finally, we translated non-English posts into English using the R package “polyglotr” to access the Google Translate API (Iwan 2023).

### 2.4. Extraction of wildlife species from Mastodon and Bluesky images

A growing focus among environmental scientists is gaining access to high-quality data on flora and fauna to support species monitoring, investigate cultural preferences for biodiversity, and create annotated imagery for training computer vision models capable of accurate species identification at scale. To evaluate whether decentralized SM platforms can provide an additional useful source of such data, we analyzed the content of the 148,285 images using the computer vision model BioCLIP (Stevens et al. 2024). The vision model is derived from the CLIP (Contrastive Language–Image Pre-training) model developed by OpenAI and trained on the TREEOFLIFE-10M dataset containing over 10 million images covering 454 thousand taxa (Van Horn and Mac Aodha 2021; Gharaee et al. 2024; Stevens et al., 2024) and 6.6M images from the Encyclopedia of Life project (https://eol.org/) covering an additional 440 thousand taxa. This rich training dataset and the advantages of the CLIP architecture have allowed BioCLIP to reach fine-grained taxonomy classification tasks (Stevens et al. 2024). BioCLIP has previously been used to detect wildlife species in social media images at multiple taxonomic levels (Klaus et al. 2026) with promising results. Here, we assessed the most frequently photographed taxonomic groups by performing an analysis at the class level, and only species identified with a confidence score of 60% or greater were included (Klaus et al. 2026). We also characterized user contribution patterns to assess potential user bias and variations in sharing behavior across taxonomic groups. For each class, we calculated the number of unique users who contributed at least one image, and summarized the distribution using the median, mean, and maximum number of images shared by a single user.

To evaluate the taxonomic representativeness of our Bluesky and Mastodon wildlife image datasets, we compared the ten most-photographed wildlife taxa on each platform with observation counts for the same taxa on iNaturalist. iNaturalist was used as an independent reference to assess whether the taxonomic composition observed on these decentralized SM was broadly consistent with patterns of biodiversity observations captured in an established citizen-science platform widely used in environmental research (Mason et al. 2025). iNaturalist counts were retrieved via its API on August 25, 2026. Cross-platform agreement was assessed using Spearman’s rank and Pearson’s correlation coefficients for each pair of comparisons.

### 2.5. Analyzing climate change discourse in Mastodon and Bluesky posts

To analyze climate change discourses on Bluesky and Mastodon, we conducted a text classification analysis using the Augmented CARDS model (Rojas et al. 2024). It is derived from the CARDS (Computer Assisted Recognition of Denial & Skepticism) model (Coan et al. 2021), which employs supervised machine learning techniques and is trained on English text from contrarian blog posts and conservative think-tank articles (Coan et al. 2021). Augmented CARDS enhances the performance of the original CARDS model by incorporating both contrarian claims about climate change (reflecting skepticism or misinformation about the cause of climate change) and convinced claims (reflecting the scientific consensus on climate change) using a large dataset of climate change-related tweets from Twitter/X (Rojas et al. 2024). The Augmented CARDS allows first, via its binary classifier, to distinguish between contrarian and convinced claims, and second, a taxonomy classifier that categorizes contrarian claims into multiple categories (Rojas et al. 2024). We applied the binary classifier to each of the 67,082 preprocessed posts collected from Bluesky and Mastodon, assigning each post to either the contrarian or convinced category. We then analyzed how the prevalence of contrarian and convinced climate change discourses changed over time across the two platforms.

## 3. Results

### 3.1. Bluesky and Mastodon in the growing decentralized SM ecosystem

Our results show clear growth in the amount and type of decentralized SM platforms after 2010 (Fig. 1). While traditional centralized mainstream platforms continue to dominate the SM landscape and maintain massive user bases, decentralized alternatives are increasingly filling similar content niches, attracting comparatively large and growing communities of users. As of June 2026, more than ten decentralized social media platforms with over 5000 users operate on the ActivityPub protocol, covering all major social media types. Mastodon remains the largest in the ecosystem, followed by the microblogging service Misskey (https://fedidb.com/software/misskey). The photo-sharing platform PixelFed, launched in 2018, has begun to grow in popularity (https://fedidb.com/software/pixelfed), while Loops, a micro-video platform, is the most recently launched platform using the ActivityPub protocol (https://joinloops.org/). By comparison, the AT Protocol ecosystem remains relatively small, consisting primarily of Bluesky and Skylight Social, a newer micro-video platform which allows users to sign in with their Bluesky account (https://skylight.social/).

The growth of user communities on the most widely adopted decentralized SM platforms suggests increasing potential for these platforms to serve as valuable data sources. Bluesky, which launched in February 2023, experienced rapid growth following the end of its invitation-only phase in July 2023. At that time, the platform had just over 200,000 users; by end of June 2026, its user base had surpassed 45 million (Fig. 2). This expansion was accompanied by a substantial increase in user engagement, with the number of unique weekly posters rising from approximately 100,000 to a peak of more than 3 million in late 2024, before settling to roughly 1.5 million by June 2026. In contrast, Mastodon, launched in 2016, has exhibited slower but steadier growth. As the largest Fediverse platform, Mastodon accounts for 69.4% of users across Fediverse platforms (https://fedidb.com/). Total users increased from approximately 7 million in July 2023 to more than 10.3 million by June 2026. Weekly active users remained relatively stable at around 1 million, following an initial peak in August 2023 and a gradual decline thereafter (Fig. 2).

**Figure 2.**
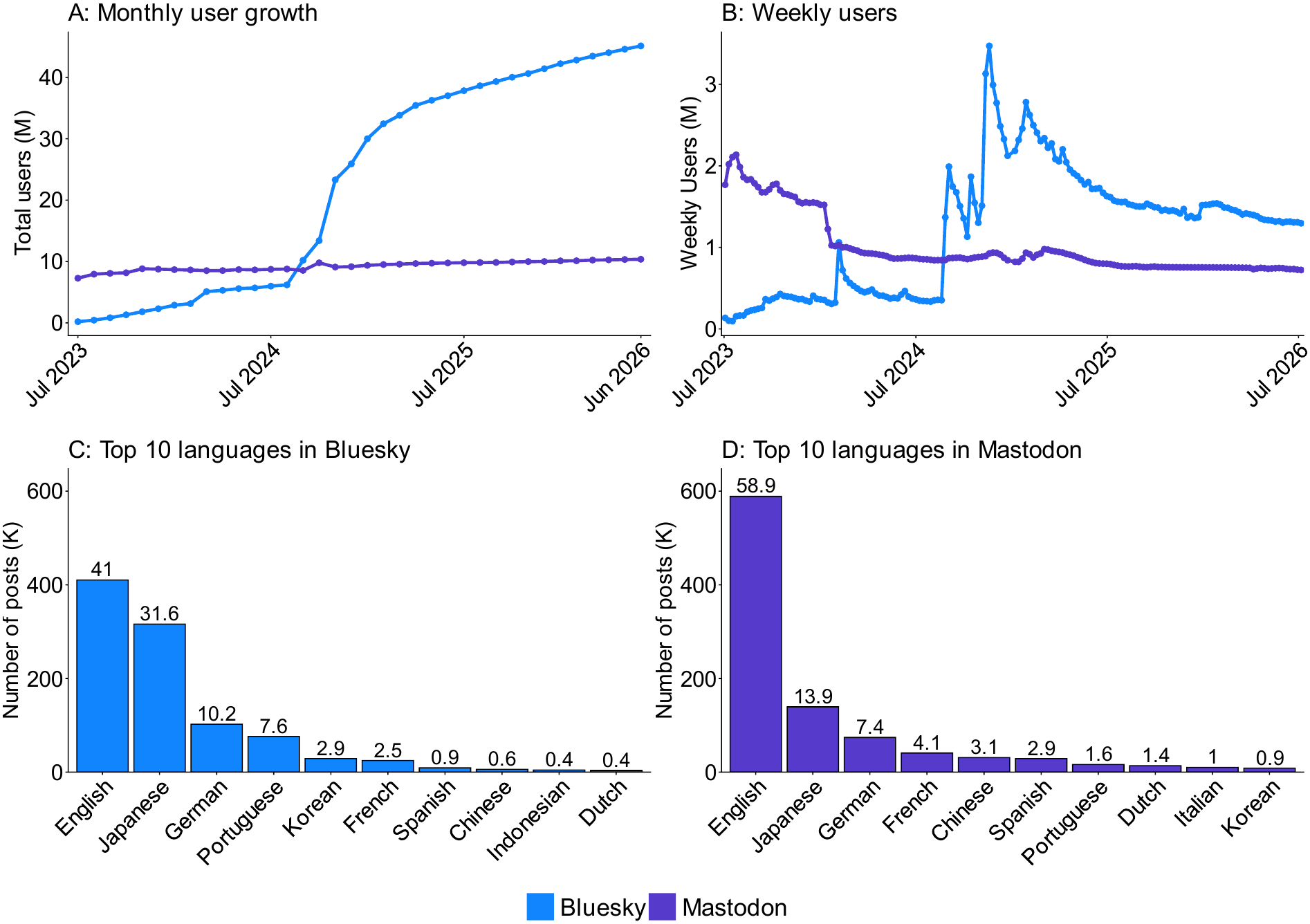
Bluesky and Mastodon users and languages. Source: User data obtained via platform APIs; language proportions computed from a dataset of 1 million randomly sampled posts.

Moreover, the linguistic diversity of user-generated content suggests broad cultural representation across the platforms, with posts appearing in 74 languages on Bluesky and 124 languages on Mastodon. While global languages still dominant with English being the most common (Bluesky: 410,000 posts, 42% of the total; Mastodon: 588,000 posts representing 58.8%), followed by Japanese (Bluesky: 316,000 31.6%; Mastodon: 139,000 posts) and German (Bluesky: 102,000, 10.2%; Mastodon: 74,000 posts), the prominence of Japanese, Chinese, and Korean among the ten most frequently used languages suggests growing adoption and engagement in East Asian countries (Fig. 2).

### 3.2. Wildlife observations in Bluesky and Mastodon

Our results revealed that Bluesky and Mastodon can expand the coverage of biodiversity assessments and study human-wildlife interactions. Content image analysis showed that Bluesky featured 116 unique species classes, while Mastodon featured 91 (the complete list can be found in Table S2 in supplementary material). Flowering plants (*Magnoliopsida*) were the most represented group, accounting for 24,479 images on Bluesky (>40% of all images) and 15,253 images on Mastodon (35%). Birds *(Aves*) were the second most photographed group, accounting for 28% of the Bluesky images and 32% of the Mastodon images. Insects (*Insecta*) and mammals (*Mammalia*) followed, each representing 16.5% and 7% of total images across both platforms. Other commonly observed wildlife (each representing less than 2% of the total) includes fungi (*Agaricomycetes*), reptiles (*Reptilia*), and amphibians (*Amphibia*), all of which appear in the top 10 most photographed species groups (Fig. 3).

**Figure 3.**
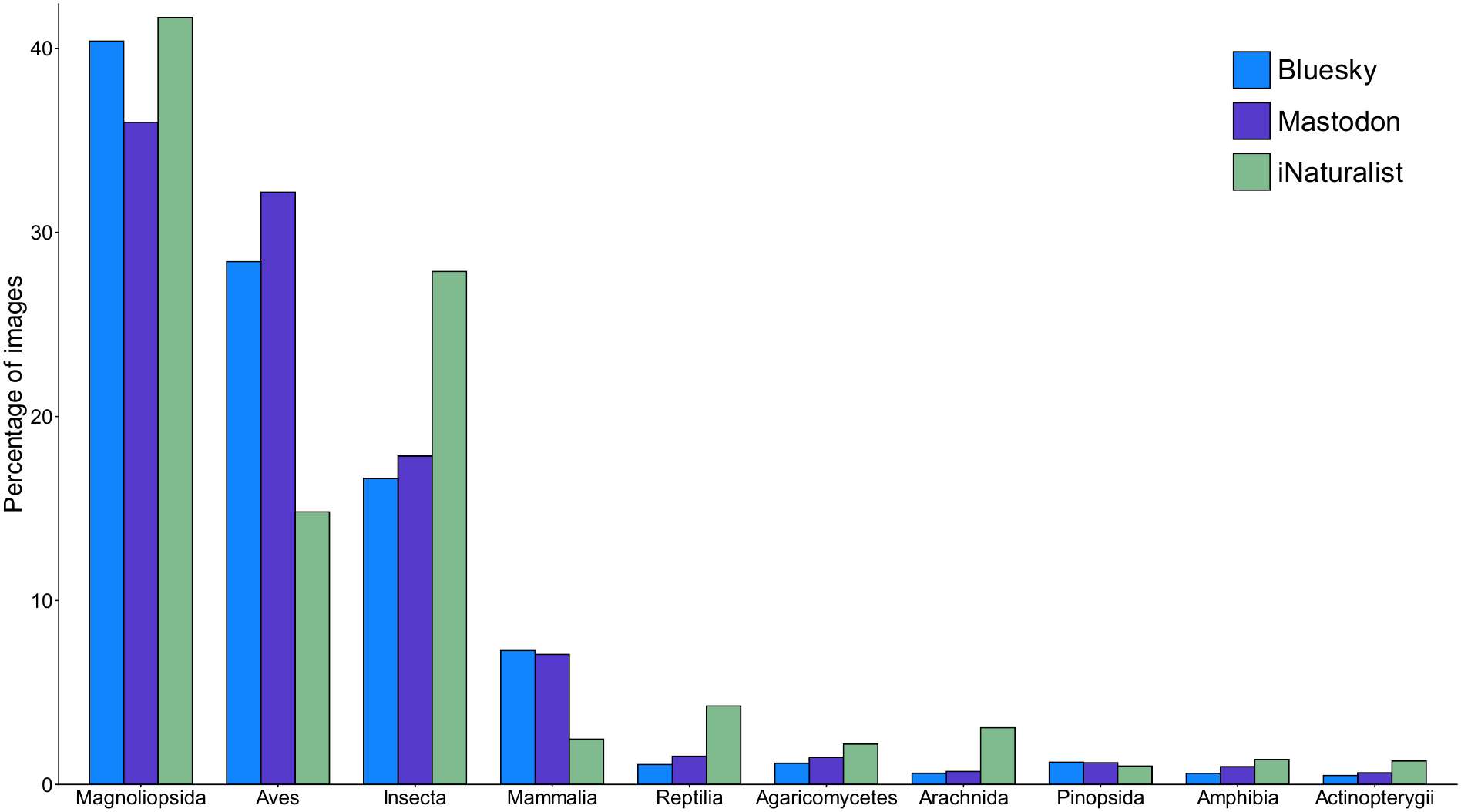
The most photographed taxa in Bluesky and Mastodon and their representation on iNaturalist. The dataset includes 60,601 images (14,242 users) on Bluesky and 42,391 images (3,179 users) on Mastodon.

Taxonomic composition on both Bluesky and Mastodon was significantly correlated with the distribution of iNaturalist observations (Bluesky: Spearman’s ρ = 0.72, p = .024, Pearson’s r = 0.90, p < .001; Mastodon: Spearman’s ρ = 0.76, p = .016, Pearson’s r = 0.87, p = .001), indicating broad consistency in the relative representation of taxa. However, compared to iNaturalist, Bluesky, and Mastodon users posted more photos of large and charismatic species (i.e., birds and mammals) and fewer of insects, fungi, and spiders.

User contribution patterns differed between Bluesky and Mastodon (Table 1). Mastodon users shared three times more images per user than Bluesky users and showed consistently higher participation across all taxonomic groups. Flowering plants (*Magnoliopsida*) had the largest contributor base on both platforms, accounting for 57.6% of all contributing users on Bluesky and 61.7% on Mastodon. Birds (*Aves*) had the highest average number of images per user on both platforms, whereas insects (*Insecta*) observations drew from a broader user base. The median contribution per user was low, at one to two images, indicating that most users shared only a small number of wildlife observations.

**Table 1.** User contribution patterns across taxonomic classes on Bluesky and Mastodon. The dataset includes 60,601 images on Bluesky and 42,391 images on Mastodon.

|  | **Bluesky** |  |  |  | **Mastodon** |  |  |  |
|---|---|---|---|---|---|---|---|---|
| **Taxonomic class** | Contributing users | Median images/ user | Mean images /user | Max images/ user | Contributing users | Median images/ user | Mean images/ user | Max images /user |
| **Magnoliopsida** | 8206 | 1 | 2.98 | 247 | 1960 | 2 | 7.67 | 681 |
| **Aves** | 3200 | 1 | 5.38 | 387 | 915 | 2 | 14.67 | 650 |
| **Insecta** | 3660 | 1 | 2.75 | 203 | 1137 | 1 | 6.47 | 571 |
| **Mammalia** | 2394 | 1 | 1.84 | 133 | 668 | 1 | 4.46 | 386 |
| **Agaricomycetes** | 342 | 1 | 2.04 | 111 | 189 | 1 | 3.26 | 38 |
| **Reptilia** | 428 | 1 | 1.53 | 21 | 181 | 1 | 3.57 | 138 |
| **Pinopsida** | 466 | 1 | 1.57 | 28 | 202 | 1 | 2.42 | 49 |
| **Amphibia** | 236 | 1 | 1.53 | 11 | 94 | 1 | 4.33 | 166 |
| **Arachnida** | 235 | 1 | 1.56 | 27 | 107 | 1 | 2.77 | 29 |
| **Actinopterygii** | 249 | 1 | 1.18 | 7 | 94 | 1 | 2.81 | 139 |
| **Other** | 878 | 1 | 1.49 | 82 | 359 | 1 | 2.45 | 152 |
| **Total** | 14,242 | 1 | 4.26 | 509 | 3179 | 2 | 13.33 | 1675 |

### 3.3. Climate change discourses in Bluesky and Mastodon

Results of analysis of climate change discussions on these platforms revealed widespread acceptance of climate change on both platforms (Fig. 4). On Bluesky, acceptance of climate change was evident in 96.9% of climate related posts (n = 31,134) compared to 3.1% *contrarian* claims (n = 997). Mastodon showed a similar distribution, with *convinced* claims comprising 97.1% of posts (n = 29,140) and *contrarian* claims accounting for 2.9% (n = 887). We also found that despite the overall increase in climate change–related posting activity, the discourse did not shift in nature and remained dominated by *convinced* claims across the entire study period.

**Figure 4.**
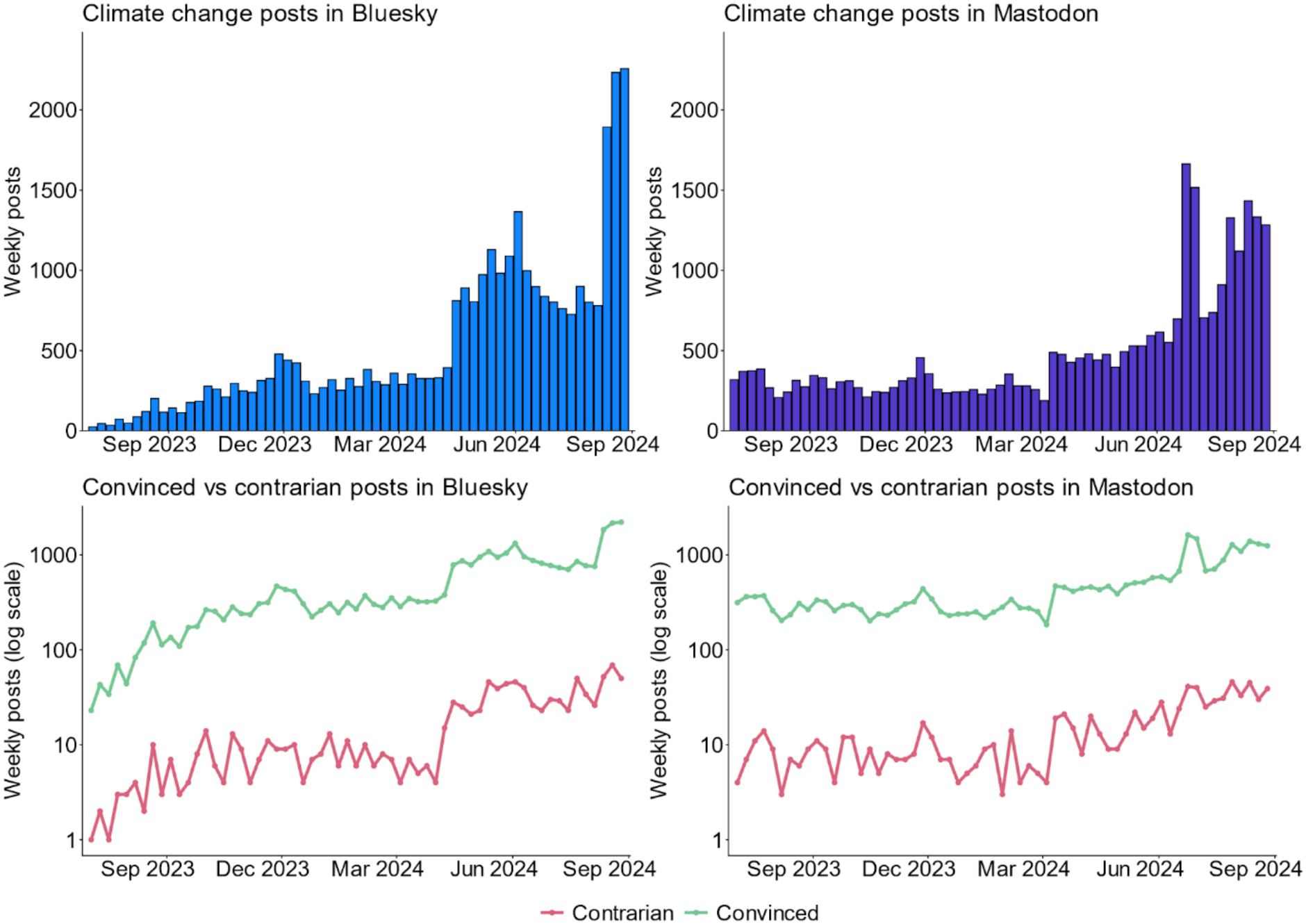
Climate change discourse on Bluesky and Mastodon: volume and stance (contrarian vs. convinced).

## 4. Discussion

Social media analysis has reshaped environmental and sustainability science (Toivonen et al. 2019; Ghermandi and Sinclair 2019; Ghermandi et al. 2023). In this paper, we analyzed emerging new social media (SM), their specificities, and evaluated their utility for environmental research through analysis of crowdsourced wildlife photography and public discourse on climate change.

Our mapping analysis revealed a proliferation of decentralized SM platforms, covering all major SM types (e.g., photo-sharing, microblogging), with new platforms launching almost annually (Fig. 1). This emergence signals a broader transition and gradual reorganization of the SM landscape (Törnberg 2026). The rise of these new alternative platforms, with millions of users migrating to them from centralized platforms such as Twitter/X and Facebook (La Cava et al. 2023; Jeong et al. 2024), is a result of increasing unease over the centralized governance of user data under a single corporation and the opacity of their algorithms (Kleppmann et al. 2024; Jeong et al. 2024), as well as recent changes in ownership and policy within the mainstream SM sphere, such as the reduction of content moderation on Twitter/X and the termination of Facebook’s fact-checking program (Jeong et al. 2024; Sabo et al. 2024). As an alternative, Bluesky, Mastodon, PixelFed, PeerTube, and other similar platforms offer a different philosophical and technical foundation, providing users with greater control over their data and more transparency in how ActivityPub and the AT Protocol operate (La Cava et al. 2022; Kleppmann et al. 2024). Among these emerging platforms, Bluesky and Mastodon are the two leading alternatives attracting the largest number of users. Our results reveal that both platforms, particularly Bluesky, have experienced sustained growth in user accounts and active participation. This upward trend suggests that they are increasingly being adopted as trusted “cyber hometowns” by users seeking alternatives to the mainstream SM ecosystem (Jeong et al. 2024; Sabo et al. 2024). Our analysis of language usage across both platforms revealed substantial linguistic diversity, with English emerging as the most prevalent language. This dominance suggests a potential spatial bias toward users in the Global North and underscores the importance of monitoring how content distribution evolves over time. Notably, we observed a significant cluster of Japanese-language content (cf. Nogara et al. 2026) and found that 3% of the Mastodon sample consisted of Chinese-language posts. This highlights the potential of decentralized platforms to reach user communities that are otherwise inaccessible through mainstream platforms such as Twitter/X, which are less popular or face regulatory restrictions in regions such as China (Pan 2017).

Despite a rapidly growing body of literature demonstrating the value of mining textual and image content from SM to monitor biodiversity and analyze climate change discourse (Ghermandi et al. 2023; Schirpke et al. 2023), recent studies have highlighted the precarious nature of relying on centralized platforms for this purpose (Ghermandi et al. 2023; Ghermandi et al. 2026). Many of these platforms have progressively restricted access to their data, limiting opportunities for environmental researchers, while Flickr, a platform that is widely used by researchers due to its consistent policy of providing fairly open access to its data, is itself experiencing a decline in popularity (Leppämäki et al. 2025). Our findings indicate that decentralized platforms offer new opportunities to revitalize SM-based environmental research, providing valuable complements—or even replacements—to extend this established body of work onto community-governed infrastructure better suited to the long-term, continuous monitoring that biodiversity and climate change demand.

By combining Bluesky and Mastodon images with the BioCLIP classifier, we demonstrated that the content of these two platforms can be used to expand the coverage of biodiversity assessments and to study people’s cultural preferences for wildlife species. Our findings reveal similar trends across both platforms: flowering plants, birds, and insects are the most frequently photographed species. This confirms previously observed trends across multiple centralized SM platforms and on iNaturalist showing that the species people share online most often are those that are common, easier to photograph, and considered more photogenic or charismatic (August et al. 2020; Havinga et al. 2023; Chai-allah et al. 2025; Klaus et al. 2026). More broadly, the diversity of species photographed on both platforms highlights their appeal to users with a range of interests in biodiversity. Moreover, our analysis of climate change posts on both platforms revealed a dominance of convinced claims over contrarian ones. This finding highlights a notable divergence between these decentralized platforms and centralized SM, particularly Twitter/X, where previous research has identified the platform as a significant vector for the dissemination of climate change skepticism and denial (Gounaridis and Newell 2024; Rojas et al. 2024). For example, Rojas et al. (2024) analyzed over 5 million climate-related tweets and found that contrarian content accounted for a daily average of 15.5%, while Gounaridis and Newell (2024) reported that 14.8% of Americans expressed climate change denial on Twitter/X, with such discourse being more prominent in conservative U.S. states. This divergence on climate discourses on Bluesky and Mastodon compared to Twitter/X, is likely attributable to the weakening content moderation policies of Twitter/X. Decentralized platforms have also attracted users migrating from Twitter/X (Jeong et al. 2024), potentially due to dissatisfaction with its policies, further reinforcing the prevalence of pro-science and climate-aware discourse within these communities. These patterns, however, may also suggest that users engaging in climate-related discussions on these platforms may differ from the broader population.

Mastodon and Bluesky open promising avenues for environmental research and beyond, due to their open protocols and APIs, which will secure more continuous access to user-generated data for scientists. However, questions remain about their adoption, for instance transitioning to a new platform involves practical and psychological efforts, particularly when it involves breaking established habits (Wood and Rünger 2016). In the case of decentralized platforms operating under ActivityPub, users must choose a server and comply with its rules. This process differs from the familiar experience of mainstream SM and may limit the broader adoption of these platforms (Wahner et al. 2024). Due to its different architecture, Bluesky’s decentralization is not directly exposed to users, which creates an experience that closely resembles that of Twitter/X (Kleppmann et al. 2024). This resemblance, combined with the inertia of the centralized platforms that currently dominate the market (Mindel et al. 2018), may partially explain why Bluesky is growing faster than Mastodon. Furthermore, the presence of numerous decentralized platforms operating under the ActivityPub protocol (Fig. 1) intensifies the competition for user attention. For instance, Mastodon, Misskey, and Pleroma are microblogging platforms that compete in a space already dominated by Twitter/X. While this diversity enables users to participate in niche communities centered on shared interests or values, it also leads to fragmentation. Finally, the user empowerment enabled by the decentralized protocols of Bluesky and Mastodon also brings challenges that scientists must consider. For example, Bluesky allows users to automatically delete older posts, while Mastodon is highly dynamic, with some servers disappearing as new ones emerge, along with shifting rules around data access (Frost-Arnold 2024; Wahner et al. 2024). Understanding how these platforms operate and engaging directly with platform communities and server administrators will be essential for maintaining access to data, gaining users’ trust in the academic use of their data and its societal value. Overall, key uncertainties and areas for inquiry for decentralized platforms remain concerning sustainable monetization models, governance frameworks, content moderation, and system scalability (Ciriello et al. 2026).

Several limitations warrant caution in interpreting our findings. First, both platforms remain smaller and demographically narrower than mainstream SM, with user bases likely skewed toward technologically engaged, English-speaking populations; the patterns we observed in wildlife photography and climate discourse may therefore not generalize to broader public attitudes. Second, our climate change classification relied on machine translation (Google Translate, via the R package polyglotr) to render non-English posts into English before applying the Augmented CARDS model, which was trained on English-language text; translation errors or loss of nuance could affect classification accuracy for non-English posts, and we did not independently validate translation quality for this dataset. Third, our species identifications from BioCLIP were retained only above a 60% confidence threshold, a choice that reduces false positives but may also exclude correctly identified but visually ambiguous or less common taxa, potentially biasing our taxonomic results toward easily distinguishable species. Finally, while we report platform growth through June 2026, our content analyses of wildlife imagery and climate discourse are based on a single collection window (January 2023–August 2024) and reflect a snapshot of two platforms that are still evolving rapidly in user composition, scale, and moderation policy; the representativeness and content patterns identified here should be re-evaluated as these platforms mature. Future research could focus on expanding the use of these new sources to other environmental and sustainability challenges, exploring their potential for finer-scale analyses at the country or regional level, and comparing their content to centralized SM.

## 5. Conclusion

This study provides one of the first systematic assessments of decentralized SM platforms, Mastodon and Bluesky, as data sources for environmental and sustainability science. Both platforms are growing steadily in users, geographic reach, and linguistic diversity, and both yield content relevant to two core sustainability domains: wildlife observation and climate change discourse. Wildlife imagery on both platforms reflects patterns long documented on centralized platforms and citizen-science tools, dominated by charismatic and easily photographed taxa such as flowering plants and birds. In contrast, climate discourse on both platforms skews heavily toward acceptance of the scientific consensus, a marked contrast to patterns reported on Twitter/X. These findings suggest that Mastodon and Bluesky can serve as viable, if imperfect, complements to mainstream platforms for environmental monitoring and public-discourse analysis, particularly as access to centralized platforms becomes more restricted and less predictable.

At the same time, our results should be read as an early, exploratory benchmark rather than a definitive characterization. Though the user bases of both platforms remain smaller and likely less demographically representative than those of mainstream platforms, they are growing and likely to evolve. The technical and governance differences between Mastodon’s ActivityPub protocol and Bluesky’s protocol-level decentralization carry distinct implications for data access and reproducibility. As these platforms continue to evolve, in scale, moderation policy, and user composition, continued monitoring will be needed to determine whether the patterns identified here persist, and whether decentralized SM can mature into a durable pillar of SM-based sustainability research rather than a transient alternative.

## Data availability

The data used in this study are open-source and publicly available. The scripts used for data collection and analysis will be made publicly available in a repository upon acceptance of the manuscript.

## References

August, T. A., O. L. Pescott, A. Joly, and P. Bonnet. 2020. AI Naturalists Might Hold the Key to Unlocking Biodiversity Data in Social Media Imagery. Patterns 1: 100116. doi:10.1016/j.patter.2020.100116.

Avalle, M., N. Di Marco, G. Etta, E. Sangiorgio, S. Alipour, A. Bonetti, L. Alvisi, A. Scala, et al. 2024. Persistent interaction patterns across social media platforms and over time. Nature 628: 582–589. doi:10.1038/s41586-024-07229-y.

Cava, L. L., L. M. Aiello, and A. Tagarelli. 2023. Drivers of social influence in the Twitter migration to Mastodon. Scientific Reports 13: 21626. doi:10.1038/s41598-023-48200-7.

Chai-allah, A., J. Hermes, A. D. La Foye, Z. S. Venter, F. Joly, G. Brunschwig, S. Bimonte, and N. Fox. 2025. Assessing recreationists’ preferences of the landscape and species using crowdsourced images and machine learning. Landscape and Urban Planning 257: 105315. doi:10.1016/j.landurbplan.2025.105315.

Ciriello, R. F., J. Marx, M. Cheong, C. Mueller-Bloch, and L. Mathiassen. 2026. Decentralized Social Media. Business & Information Systems Engineering 68: 219–234. doi:10.1007/s12599-025-00952-4.

Coan, T. G., C. Boussalis, J. Cook, and M. O. Nanko. 2021. Computer-assisted classification of contrarian claims about climate change. Scientific Reports 11: 22320. doi:10.1038/s41598-021-01714-4.

Failla, A., and G. Rossetti. 2024. “I’m in the Bluesky Tonight”: Insights from a year worth of social data. Edited by Fabio Saracco. PLOS ONE 19: e0310330. doi:10.1371/journal.pone.0310330.

Falkenberg, M., A. Galeazzi, M. Torricelli, N. Di Marco, F. Larosa, M. Sas, A. Mekacher, W. Pearce, et al. 2022. Growing polarization around climate change on social media. Nature Climate Change 12: 1114–1121. doi:10.1038/s41558-022-01527-x.

Frost-Arnold, K. 2024. Beyond Corporate Social Media Platforms: The Epistemic Promises and Perils of Alternative Social Media. Topoi 43: 1557–1568. doi:10.1007/s11245-024-10102-2.

Gharaee, Z., Z. Gong, N. Pellegrino, I. Zarubiieva, J. B. Haurum, S. C. Lowe, J. T. A. McKeown, C. C. Y. Ho, et al. 2023. A Step Towards Worldwide Biodiversity Assessment: The BIOSCAN-1M Insect Dataset (version 3). arXiv. doi:10.48550/ARXIV.2307.10455.

Ghermandi, A., and M. Sinclair. 2019. Passive crowdsourcing of social media in environmental research: A systematic map. Global Environmental Change 55: 36–47. doi:10.1016/j.gloenvcha.2019.02.003.

Ghermandi, A., J. Langemeyer, D. Van Berkel, F. Calcagni, Y. Depietri, L. Egarter Vigl, N. Fox, I. Havinga, et al. 2023. Social media data for environmental sustainability: A critical review of opportunities, threats, and ethical use. One Earth 6: 236–250. doi:10.1016/j.oneear.2023.02.008.

Ghermandi, A., F. Calcagni, J. Langemeyer, and U. Schirpke. 2026a. Digital windows into nature’s values: A critical review of cultural ecosystem services research with social media data. Ecosystem Services 79: 101839. doi:10.1016/j.ecoser.2026.101839.

Ghermandi, A., Y. Depietri, and D. E. Orenstein. 2026b. Human-nature interactions through a digital prism: Heterogeneity in user socio-demographics and content across social media platforms. Landscape and Urban Planning 268: 105568. doi:10.1016/j.landurbplan.2025.105568.

Gounaridis, D., and J. P. Newell. 2024. The social anatomy of climate change denial in the United States. Scientific Reports 14: 2097. doi:10.1038/s41598-023-50591-6.

Guess, A. M., N. Malhotra, J. Pan, P. Barberá, H. Allcott, T. Brown, A. Crespo-Tenorio, D. Dimmery, et al. 2023. How do social media feed algorithms affect attitudes and behavior in an election campaign? Science 381: 398–404. doi:10.1126/science.abp9364.

Hanny, D., S. Schmidt, and B. Resch. 2026. Bluesky as a social media data source for disaster management: investigating spatio-temporal, semantic and emotional patterns for floods and wildfires. Journal of Computational Social Science 9: 16. doi:10.1007/s42001-025-00448-x.

Havinga, I., D. Marcos, P. Bogaart, D. Massimino, L. Hein, and D. Tuia. 2023. Social media and deep learning reveal specific cultural preferences for biodiversity. People and Nature 5: 981–998. doi:10.1002/pan3.10466.

Jeong, D., S. Hwang, J. Kim, H. Yu, and E. Park. 2023. Public perspective on renewable and other energy resources: Evidence from social media big data and sentiment analysis. Energy Strategy Reviews 50: 101243. doi:10.1016/j.esr.2023.101243.

Jeong, U., A. Nirmal, K. Jha, S. X. Tang, H. R. Bernard, and H. Liu. 2023. User Migration across Multiple Social Media Platforms (version 2). arXiv. doi:10.48550/ARXIV.2309.12613.

Johnson, N. F., R. Leahy, N. J. Restrepo, N. Velasquez, M. Zheng, P. Manrique, P. Devkota, and S. Wuchty. 2019. Hidden resilience and adaptive dynamics of the global online hate ecology. Nature 573: 261–265. doi:10.1038/s41586-019-1494-7.

Kim, A., and A. R. Dennis. 2019. *Says Who?* The Effects of Presentation Format and Source Rating on Fake News in Social Media1. MIS Quarterly 43: 1025–1039. doi:10.25300/MISQ/2019/15188.

Kitchens, B., S. L. Johnson, and P. Gray. 2020. Understanding Echo Chambers and Filter Bubbles: The Impact of Social Media on Diversification and Partisan Shifts in News Consumption. MIS Quarterly 44: 1619–1649. doi:10.25300/MISQ/2020/16371.

Klaus, V. H., F. J. Richter, N. Fox, D. Andreatta, and A. Chai-allah. 2026. Powerful flowers: Public perception of grassland aesthetics is strongly related to management and biodiversity. Agriculture, Ecosystems & Environment 398: 110111. doi:10.1016/j.agee.2025.110111.

Kleppmann, M., P. Frazee, J. Gold, J. Graber, D. Holmgren, D. Ivy, J. Johnson, B. Newbold, et al. 2024. Bluesky and the AT Protocol: Usable Decentralized Social Media (version 2). arXiv. doi:10.48550/ARXIV.2402.03239.

La Cava, L., S. Greco, and A. Tagarelli. 2022. Information consumption and boundary spanning in Decentralized Online Social Networks: The case of Mastodon users. Online Social Networks and Media 30: 100220. doi:10.1016/j.osnem.2022.100220.

Leppämäki, T., V. Heikinheimo, J. Eklund, A. Hausmann, and T. Toivonen. 2025. The rise and fall of the social media platform Flickr: Implications for nature recreation research. Journal of Outdoor Recreation and Tourism 50: 100880. doi:10.1016/j.jort.2025.100880.

Ma, B., and R. Zheng. 2025. Exploring Food Safety Emergency Incidents on Sina Weibo: Using Text Mining and Sentiment Evolution. Journal of Food Protection 88: 100418. doi:10.1016/j.jfp.2024.100418.

Mason, B. M., T. Mesaglio, J. Barratt Heitmann, M. Chandler, S. Chowdhury, S. B. Z. Gorta, F. Grattarola, Q. Groom, et al. 2025. iNaturalist accelerates biodiversity research. BioScience 75: 953–965. doi:10.1093/biosci/biaf104.

Mindel, V., L. Mathiassen, and A. Rai. 2018. The Sustainability of Polycentric Information Commons1. MIS Quarterly 42: 607–631. doi:10.25300/MISQ/2018/14015.

Moore, F. C., and N. Obradovich. 2020. Using remarkability to define coastal flooding thresholds. Nature Communications 11: 530. doi:10.1038/s41467-019-13935-3.

Nogara, G., E. S. Sahneh, M. R. DeVerna, N. Liu, L. Luceri, F. Menczer, F. Pierri, and S. Giordano. 2026. A longitudinal analysis of misinformation, polarization and toxicity on Bluesky after its public launch. Online Social Networks and Media 51: 100342. doi:10.1016/j.osnem.2025.100342.

Pan, J. 2017. How Market Dynamics of Domestic and Foreign Social Media Firms Shape Strategies of Internet Censorship. Problems of Post-Communism 64: 167–188. doi:10.1080/10758216.2016.1181525.

Rojas, C., F. Algra-Maschio, M. Andrejevic, T. Coan, J. Cook, and Y.-F. Li. 2024. Hierarchical machine learning models can identify stimuli of climate change misinformation on social media. Communications Earth & Environment 5: 436. doi:10.1038/s43247-024-01573-7.

Roscam Abbing, R., C. Diehm, and S. Warreth. 2023. Decentralised social media. Internet Policy Review 12. doi:10.14763/2023.1.1681.

Sabo, E., T. Gesthuizen, K. J. A. Bouma, D. Karastoyanova, and M. Riveni. 2024. An analysis of mastodon adoption dynamics based on instance types. Social Network Analysis and Mining 14: 184. doi:10.1007/s13278-024-01341-7.

Schirpke, U., A. Ghermandi, M. Sinclair, D. Van Berkel, N. Fox, L. Vargas, and L. Willemen. 2023. Emerging technologies for assessing ecosystem services: A synthesis of opportunities and challenges. Ecosystem Services 63: 101558. doi:10.1016/j.ecoser.2023.101558.

Shiffman, D. 2025. Social media for fisheries science and management professionals: How to use Bluesky and Instagram, and why you should. Fisheries 50: 316–322. doi:10.1093/fshmag/vuaf025.

Stevens, S., J. Wu, M. J. Thompson, E. G. Campolongo, C. H. Song, D. E. Carlyn, L. Dong, W. M. Dahdul, et al. 2024. BioCLIP: A Vision Foundation Model for the Tree of Life. In 2024 IEEE/CVF Conference on Computer Vision and Pattern Recognition (CVPR), 19412–19424. Seattle, WA, USA: IEEE. doi:10.1109/CVPR52733.2024.01836.

Toivonen, T., V. Heikinheimo, C. Fink, A. Hausmann, T. Hiippala, O. Järv, H. Tenkanen, and E. Di Minin. 2019. Social media data for conservation science: A methodological overview. Biological Conservation 233: 298–315. doi:10.1016/j.biocon.2019.01.023.

Törnberg, P. 2026. Shifts in U.S. Social Media Use, 2020–2024: Decline, Fragmentation, and Enduring Polarization. Journal of Quantitative Description: Digital Media 6. doi:10.51685/jqd.2026.005.

Van Zanten, B. T., D. B. Van Berkel, R. K. Meentemeyer, J. W. Smith, K. F. Tieskens, and P. H. Verburg. 2016. Continental-scale quantification of landscape values using social media data. Proceedings of the National Academy of Sciences 113: 12974–12979. doi:10.1073/pnas.1614158113.

Wähner, M., A. Deubel, J. Breuer, and K. Weller. 2024. “Don’t research us”—How Mastodon instance rules connect to research ethics. Publizistik 69: 357–380. doi:10.1007/s11616-024-00855-6.

Williams, H. T. P., J. R. McMurray, T. Kurz, and F. Hugo Lambert. 2015. Network analysis reveals open forums and echo chambers in social media discussions of climate change. Global Environmental Change 32: 126–138. doi:10.1016/j.gloenvcha.2015.03.006.

Wood, W., and D. Rünger. 2016. Psychology of Habit. Annual Review of Psychology 67: 289–314. doi:10.1146/annurev-psych-122414-033417.

Zheng, E.-T., X. Jiang, Z. Fang, and M. Thelwall. 2025. How is science discussed on Bluesky? (version 2). arXiv. doi:10.48550/ARXIV.2507.18840.

Zignani, M., S. Gaito, and G. P. Rossi. 2018. Follow the “Mastodon”: Structure and Evolution of a Decentralized Online Social Network. Proceedings of the International AAAI Conference on Web and Social Media 12: 541–550. doi:10.1609/icwsm.v12i1.14988.

Zulli, D., M. Liu, and R. Gehl. 2020. Rethinking the “social” in “social media”: Insights into topology, abstraction, and scale on the Mastodon social network. New Media & Society 22: 1188–1205. doi:10.1177/1461444820912533.

